# Mathematical modeling of the effects of Wnt-10b on bone metabolism

**DOI:** 10.1101/2021.06.12.448204

**Authors:** Carley V. Cook, Mohammad Aminul Islam, Brenda J. Smith, Ashlee N. Ford Versypt

**Author notes:** **Correspondence** Ashlee N. Ford Versypt, Department of Chemical and Biological Engineering, University at Buffalo, The State University of New York, 507 Furnas Hall, Buffalo, NY, 14260, USA. **Funding information** National Institutes of Health, Grant/Award Number: R35GM133763.

## Abstract

Bone health is determined by factors including bone metabolism or remodeling. Wnt-10b alters osteoblastogenesis through pre-osteoblast proliferation and differentiation and osteoblast apoptosis rate, which collectively lead to the increase of bone density. To model this, we adapted a previously published model of bone remodeling. The resulting model for the bone compartment includes differential equations for active osteoclasts, pre-osteoblasts, osteoblasts, osteocytes, and the amount of bone present at the remodeling site. Our alterations to the original model consist of extending it past a single remodeling cycle and implementing a direct relationship to Wnt-10b. Four new parameters were estimated and validated using normalized data from mice. The model connects Wnt-10b to bone metabolism and predicts the change in trabecular bone volume caused by a change in Wnt-10b input. We find that this model predicts the expected increase in pre-osteoblasts and osteoblasts while also pointing to a decrease in osteoclasts when Wnt-10b is increased.

## Introduction

Osteoporosis, characterized by decreased bone mass and structural deterioration, results from an imbalance in the tissue’s metabolic processes. In the adult skeleton, bone is remodeled within millions of basic multicellular units (BMU) and involves the activity of osteoclasts, osteoblasts, osteocytes, and their precursors^1–4^. An important regulator in bone remodeling is the canonical Wnt/*β*-catenin pathway^5,6^. This pathway is important for growth and development^6^, maintenance of teeth^6^, and immune responses^7–9^. This pathway is activated by glycoproteins in the wingless-related integration site family (Wnt). Of the nineteen different types of Wnt, only seven are known to activate the canonical Wnt/*β*-catenin pathway. Activation with a fizzled receptor and a low-density lipoprotein receptor-related protein coreceptor bound to Wnt prevents *β*-catenin deconstruction by the proteasome.^6,8^. Avoiding deconstruction, *β*-catenin activates gene expressions related to Wnt expression in the nucleus.

Expression of Wnt-10b, a canonical Wnt/*β*-catenin pathway activator, occurs in various systems in the body including both the bone microenvironment and the immune system^10^. While over activation of the canonical Wnt/*β*-catenin pathway has been associated with a tumor environment ^7,9,10^, normal levels of Wnt-10b are required for T cells to develop and function properly ^10^. When targets of the *β*-catenin pathway are disrupted, T cells do not properly develop. In one study, T cell factor-1 was disrupted in a mouse model causing the T cells to not develop properly^11^, and in another study complete deletion of lymphoid enhancing factor-1 led to nonviable mice^12^. T cells can also be sources of Wnt-10b^10,13^.

T-cell produced Wnt-10b has been implicated in altering bone volume in mice^13,14^ and has been simulated by our team^15^. This suggests another link between the bone and the immune system as Wnt-10b is necessary for bone remodeling. Wnt-10b can also be produced by cells of osteoblastic lineage. A change in Wnt-10b levels has been shown to alter bone volume significantly ^13,16–18^. Wnt-10b regulates bone volume via the canonical pathway by activating genes related to osteoblastogenesis, pre-osteoblast proliferation, and cell survival ^10,19,20^. The extent to which changes in Wnt-10b levels alter these factors and how these alterations affect the cell populations and bone volume as a whole is currently unknown.

Currently, there are a few published mathematical/computational models on bone metabolism. One model describes the autocrine and paracrine interactions of osteoblasts and osteoclasts using ordinary differential equations (ODEs) that track the cell populations as well as changes in trabecular bone mass^21^. This model has been updated to include a relationship with the parathyroid hormone (PTH)^22^. These publications along with a model that tracks spatial trabecular bone volume changes ^23^ have been used to develop a human based ODE model that tracks how different signals cause the development of active osteoclasts and osteoblasts and a change in trabecular bone mass^24^. None of these models include Wnt-10b; therefore, each lacks a vital part of the bone remodeling system. Proctor and Garland ^25^ have pointed out the importance of including Wnt-10b in a bone remodeling model using a network model that identifies the Wnt and PTH pathways as the most important systems in regulating bone remodeling.

A few of the existing bone metabolism models do explore the role Wnt-10b has in bone remodeling^26–30^. However, the models in one subset ^26–28^ assume a constant basal concentration of Wnt-10b and focus on how an increase of sclerostin competes for the Wnt binding sites. The models in the other subset^29,30^ allow the Wnt-10b concentration to vary but only in the presence of a tumor. These models connect Wnt-10b through Hill functions ^31^ that alter an assortment of the terms for pre-osteoblast proliferation, differentiation from pre-osteoblasts to osteoblasts, and osteoblast apoptosis (Table 1). None of these models address potential changes in Wnt-10b levels caused by T cells or BMU cells.

**TABLE 1.** Wnt-10b relationships presented in existing models for bone remodeling.

| Model | Wnt-10b type | Wnt-10b interactions |
| --- | --- | --- |
| 26 | Constant | Not explicitly modeled |
| 27 | Constant | Pre-osteoblast proliferation |
| 28 | Constant | Pre-osteoblast proliferation<br>Pre-osteoblast differentiation<br>Osteoblast apoptosis |
| 29 | Dynamic | Pre-osteoblast proliferation |
| 30 | Dynamic | Pre-osteoblast proliferation<br>Pre-osteoblast differentiation |

The present work aims to improve understanding of bone metabolism by developing a computational model that directly connects the remodeling cycle to altered Wnt-10b concentrations that can be constant or dynamically induced by other physiologically processes such as an immune response. The model development is covered in the Methods section, which includes an overview of the model, the data used for the model, and the techniques used to determine parameter values. After validating the model, we used it to explore how Wnt-10b directly affects important cell populations in a BMU and how these population changes alter bone volume (Results and Discussion). The model will allow us in the future to explore potential ways to shift bone metabolism away from an imbalance that favors bone resorption to prevent bone loss or restore bone, but not without some limitations.

## Methods

We alter a published model, which is referred to as Graham 2013^24^, to include a mathematical relationship that represents how Wnt-10b interacts with bone formation (Figure 1). This is a single compartment model that tracks the important cells involved in the bone remodeling cycle as well as bone volume. We extend past a single remodeling cycle and add a direct relationship to Wnt-10b stimuli. We add a relationship to Wnt-10b by utilizing published data to fit model parameters.

**FIGURE 1.**
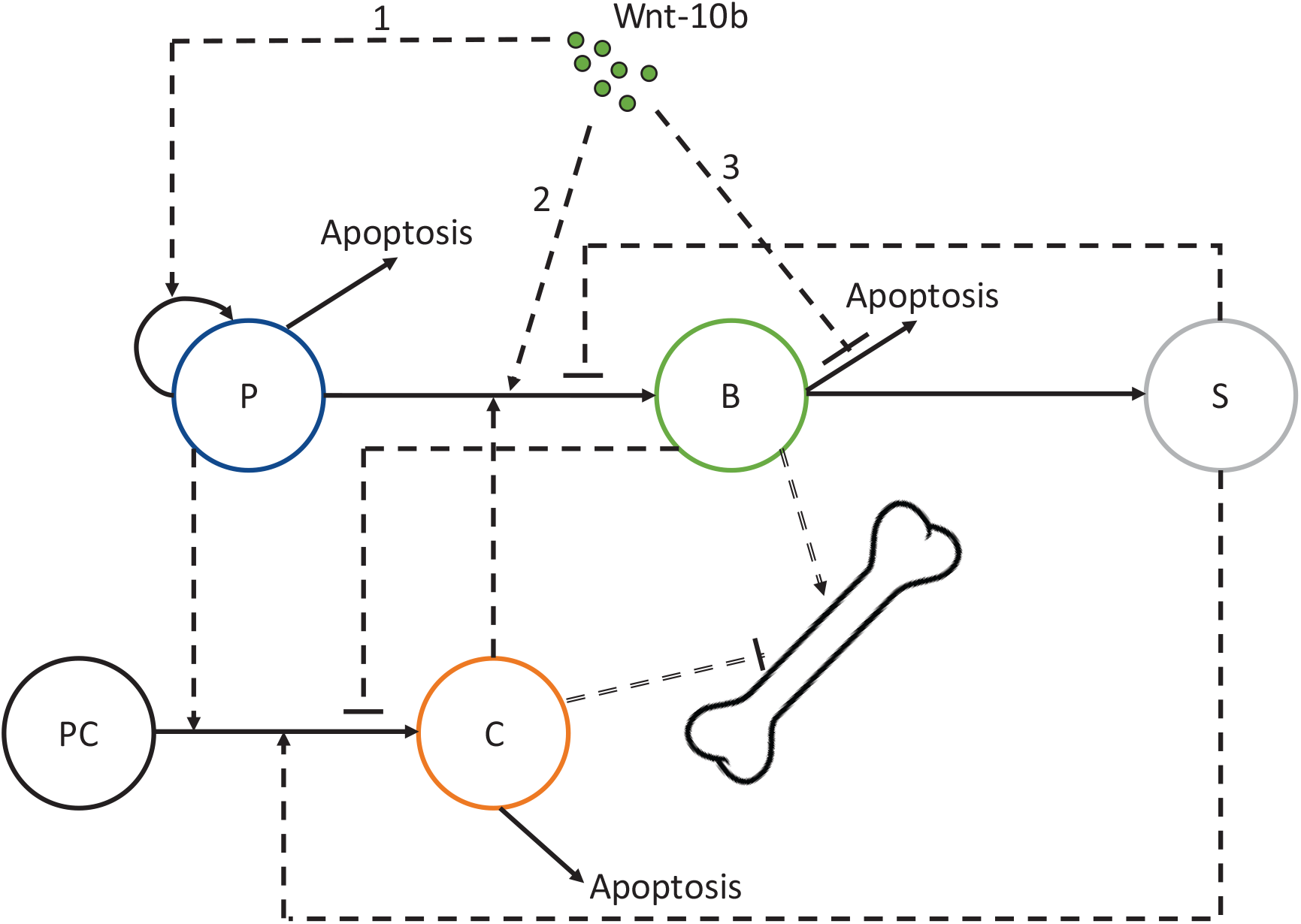
Interactions between bone cell populations considered in the Graham 2013 model^24^ and the added impact of Wnt-10b (labels 1-3). Pre-osteoblasts (P), osteoblasts (B), osteocytes (S), and osteoclasts (C) are tracked by differential equations. Pre-osteoclasts (PC) are assumed constant and are not explicitly modeled. The interactions include an assortment of autocrine (not depicted here) and paracrine (single dashed lines) signaling factors. Single dashed lines with arrows represent activating relationships, and single dashed lines with flat ends represent inhibitory relationships. Solid lines represent transformations: self-proliferation, differentiation, and apoptosis. Wnt-10b (Wnt) stimulates osteoblastogenesis through pre-osteoblast proliferation (1) and differentiation to osteoblasts (2). Wnt-10b inhibits osteoblast apoptosis (3). Impacts of bone cells on bone volume from a combination of bone resorption and formation are represented by double dashed lines with flat and arrow ends, respectively.

### Graham 2013 model

The Graham 2013 model includes five ODEs that track the changes in bone volume (*z*) through the active populations of osteocytes (*S*), pre-osteoblasts (*P*), osteoblasts (*B*), and osteoclasts (*C*)^24^. The model does not include an equation for the population of pre-osteoclasts. The model includes important autocrine and paracrine signaling factors that are represented by power law relationships, *g*_*ij*_ and *f*_*ij*_, where *i* and *j* are the two cell types involved in the signaling. This section summarizes the parts of the Graham 2013 model^24^ and parameter values that we adopt directly without alteration (Table 2).

**TABLE 2.** Parameter values and definitions directly from the Graham 2013 model^24^.

| Parameter | Definition | Value | Units |
| --- | --- | --- | --- |
| $\alpha_1$ | Osteoblast embedding rate | 0.5 | day<sup>-1</sup> |
| $\alpha_2$ | Differentiation rate of pre-osteoblast precursors | 0.1 | day<sup>-1</sup> |
| $\alpha_3$ | Pre-osteoblast proliferation rate | 0.1 | day<sup>-1</sup> |
| $\alpha_4$ | Differentiation rate of osteoclast precursors | 0.1 | day<sup>-1</sup> |
| $\beta_1$ | Differentiation rate of pre-osteoblasts | 0.1 | day<sup>-1</sup> |
| $\beta_2$ | Osteoblast apoptosis rate | 0.1 | day<sup>-1</sup> |
| $\delta$ | Pre-osteoblast apoptosis rate | 0.1 | day<sup>-1</sup> |
| $\epsilon$ | Avoid 0 denominator | 1 | cells |
| $K_s$ | Critical value of osteocyte population | 200 | cells |
| $g_{31}$ | Osteoblast autocrine signaling | 1 | dimensionless |
| $g_{21}$ | Osteocyte paracrine signaling of pre-osteoblasts | 2 | dimensionless |
| $g_{22}$ | Sclerostin regulation of osteoblastogenesis | 1 | dimensionless |
| $g_{32}$ | Pre-osteoblast autocrine signaling | 1 | dimensionless |
| $g_{41}$ | Osteocyte paracrine signaling of osteoclasts | 1 | dimensionless |
| $g_{42}$ | Pre-osteoblast paracrine signaling of osteoclasts | 1 | dimensionless |
| $g_{43}$ | Osteoblast paracrine signaling of osteoclasts | -1 | dimensionless |
| $g_{44}$ | Sclerostin regulation of osteoclastogenesis | 1 | dimensionless |
| $f_{12}$ | Pre-osteoblast paracrine signaling of osteoblasts | 1 | dimensionless |
| $f_{14}$ | Osteoclast paracrine signaling of osteoblasts | 1 | dimensionless |
| $f_{23}$ | Osteoblast autocrine signaling for apoptosis | 1 | dimensionless |
| $f_{34}$ | Osteoclast autocrine signaling for apoptosis | 1 | dimensionless |
| $k_1$ | Bone resorption rate | 0.698331 | %volume cells<sup>-1</sup> day<sup>-1</sup> |
| $k_2$ | Balanced bone formation rate | 0.015445 | %volume cells<sup>-1</sup> day<sup>-1</sup> |

Note: $k_1$ was manually adjusted from the rounded value of 0.7 reported in the Graham 2013 model<sup>24</sup> to six significant figures as for $k_2$ to maintain no steady state change in bone volume at the end of a remodeling cycle (in the absence of Wnt-10b perturbations) when the populations of all cell types return to their initial conditions.

### Osteocytes

Equation 1 describes the dynamics of the osteocyte cell population, *S*. Mature osteoblasts convert into an osteocyte phenotype at a rate of *α*_1_. The term 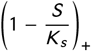 represents the effectiveness of sclerostin regulation by osteocytes, and the subscript + means that the value must remain greater than or equal to zero. If the quantity inside the parentheses were to be evaluated as less than zero, then the term would be set to zero. Note that although sclerostin regulation does include a Wnt pathway, we are focusing on Wnt-10b excreted from T cells or from a genetic perturbation, not Wnt-10b produced within a normal balanced remodeling cycle. It is assumed in the model that over the duration of remodeling, osteocytes will not die. Instead of a death term, the osteocyte population is reduced from the steady state value of 200 cells to 180 cells at the start of each remodeling cycle ^32–35^. This reduction leads to a decreased level of sclerostin and allows for the RANKL activation of osteoclasts (Equation 2). This process represents the initial biomechanical action that triggers a remodeling cycle.

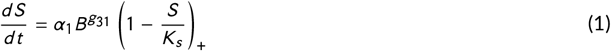

### Osteoclasts

Equation 2 describes the dynamics of the activated osteoclast cell population, *C*. A large amount of pre-osteoclasts are assumed to be available without a significant change in the population, so the pre-osteoclast population is not modeled. The production of osteoclasts depends on a differentiation rate of *α*_4_ and an interaction of the receptor activator of NF-*κ*B (RANK)/RANK ligand (RANKL)/osteoprotegerin (OPG) system that is described by the term 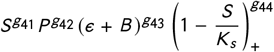. OPG from osteoblasts can act as a decoy receptor for RANKL. This interaction is represented as (*ϵ* + *B*)^*g*43^. This term includes a very small number, *ϵ*, to prevent dividing by zero when the osteoblast population is zero, since *g*_43_ is a negative integer. The second part of the equation represents osteoclast apoptosis with a first order rate constant *β*_3_.

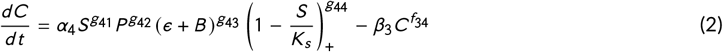

### Model enhancements for Wnt-10b dependence

This section covers the parts of the model that we alter to explicitly account for Wnt-10b changes from stimuli outside the normal bone remodeling cycle. Several parts of the equations remain the same as the Graham 2013 model (Table 2), but with new terms added or parameters adjusted in each of the following equations (Table 5). The adjusted parameters are denoted with the subscript *adj*. As these additions are related to Wnt-10b, we introduce a variable *Wnt* into the equations that represents the normalized fold change of Wnt-10b present compared to the normal levels of Wnt-10b in the system. Note that if the Wnt-10b levels are normal, the remodeling cycle is normal as well, and *Wnt* takes on a value of zero. Figure 1 visually describes how Wnt-10b alters the remodeling cycle.

We connect the concentration of Wnt-10b to the remodeling cycle through a Hill function in the form of

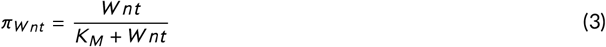

We assume that *K*_*M*_ remains the same for both the pre-osteoblast and osteoblast interactions because they are both from osteoblast lineage. A Hill function relationship is chosen based on the previous models related to Wnt-10b (Table 1). This particular form is chosen so that when the Wnt-10b concentration is not perturbed, *π*_*Wnt*_ will take the value of zero, and the model will return to a steady state of no net bone volume change at the end of a remodeling cycle.

### Pre-osteoblasts

Equation 4 describes the dynamics of the pre-osteoblast cell population, *P*. The first four terms are unchanged from the Graham 2013 model. Pre-osteoblasts differentiate at a rate of *α*_2_ from a large population of stem cells. This differentiation is triggered by the decrease in sclerostin signaling of osteocytes at the beginning of a remodeling cycle. The pre-osteoblast cell population can also be increased by proliferation of existing cells at a rate of *α*_3_. The pre-osteoblast population is decreased by differentiation into osteoblasts regulated by paracrine signals from the osteoclast population and by apoptosis with a first order rate constant of *δ*. The new terms we include here account for Wnt-10b effects on the pre-osteoblast proliferation and pre-osteoblast differentiation to osteoblasts, both via the factor defined in Equation 3. The fifth term in Equation 4 represents the increase in pre-osteoblast proliferation rate due to excess Wnt-10b. The sixth term quantifies Wnt-10b direct effects on pre-osteoblast differentiation independent from osteoclast regulation.

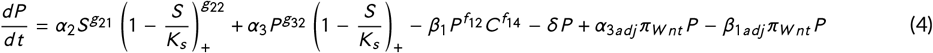

### Osteoblasts

Equation 5 describes the dynamics of the osteoblast cell population, *B*. Pre-osteoblasts mature into osteoblasts at a rate of *β*_1_ with osteoclast regulation and normal Wnt-10b (*Wnt* = 0), but when the amount of Wnt-10b is altered, the maturation rate changes by *β*_1*adj*_ *π*_*Wnt*_ as discussed above for Equation 4. Wnt-10b also changes how fast osteoblasts die. We reduce the osteoblast apoptosis rate constant *β*_2_ by *β*_2*adj*_ *π*_*Wnt*_. The final term in Equation 5 represents the conversion of osteoblasts into osteocytes.

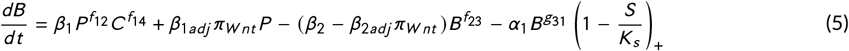

### Trabecular bone volume

Equation 6 shows the dynamics of trabecular bone volume, *z*, at a single remodeling site. Bone volume is reduced by osteoclasts at a rate of *k*_1_. This rate is slightly different than the rate in Graham 2013 due to rounding. In the Graham model, the reported *k*_1_ value results in a slightly unbalanced remodeling cycle (a new steady state results at the end of each cycle and amplified after repeated cycles). We manually adjust *k*_1_ to match the same number of significant figures as *k*_2_ until we obtain a completely balanced remodeling cycle–that is, no steady state change in bone volume at the end of one remodeling cycle (in the absence of Wnt-10b perturbations) when the populations of all cell types return to their initial conditions. Note that this alteration is not related to a change in Wnt-10b concentration. Osteoblasts build back the bone at a rate of *k*_2_.

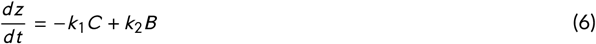

### Mouse data

This section covers the data used and the assumptions made concerning the data during the development of the model. Data originally presented in graphs was extracted using Plot Digitizer (version 2.6.8)^36^, a tool that digitizes the axes and returns the data points based on the pixel locations. We utilized three published data sources from *in vivo* experiments with C57BL/6 mice (Table 3). Two sources from the same lab ^16,17^ were used to estimate the parameters, and the third source from another lab ^18^ was used to validate the model. Calibration and validation were done using mice data for two reasons. The first is that mice data is the only type of data that we currently have access to. The second is that mice are commonly used to study age related bone loss due to the similarities in mice and human bone remodeling^37^.

**TABLE 3.** Bone volume data from mice with genetically altered Wnt-10b expression.

| Normalized<br>Wnt-10b<br>fold change | Age<br>(months) | Altered<br>BV/TV (%) | Altered SD<br>BV/TV (%) | Baseline<br>BV/TV (%) | Normalized<br>BV/TV | Source |
| --- | --- | --- | --- | --- | --- | --- |
| -1 | 2 | 6.4 | 1.8 | 9.1 | -29.7 | 17 |
| -1 | 3 | 4.3 | 1 | 7.4 | -41.9 | 16 |
| +1.8 | 3 | 8.04 | 1.22 | 6.35 | 26.6 | 18 |
| +1.8 | 6 | 4.42 | 1.32 | 3.24 | 36.6 | 18 |
| +5 | 3 | 18.1 | 4.3 | 10.7 | 69.2 | 16 |
| +50 | 6 | 15.8 | 4.1 | 3.6 | 339 | 17 |

Note: To produce the 50-fold change of Wnt-10b in mice, Bennett et al.<sup>17</sup> followed the procedures in Longo et al.<sup>43</sup>.

### Trabecular bone volume

The trabecular bone volume per total volume (BV/TV) data was normalized against the control groups using

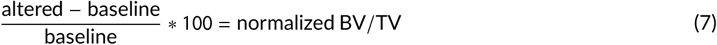

turning BV/TV data into relative change in trabecular bone volume. We refer to “altered BV/TV” to denote mice that were genetically altered to over- or under-produce Wnt-10b, while “baseline BV/TV” denotes unaltered litter mates. To visualize the most conservative amount of uncertainty in the data, we only considered the standard deviation (SD) of the altered mice. Therefore, we took the uncertainty to be

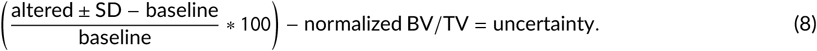

The uncertainty was used only for a visual check as this study is focused on the average of a population, not the variance within a population.

We normalized the mouse data for use in the model previously parameterized for humans. We assumed that the normalized BV/TV data is consistent from mice to human. Note that since our model starts at a baseline of 100 percent of normal bone volume, we expect our simulation to produce *z* that is 100 plus the expected normalized bone volume. We also took into account the different remodeling cycle lengths of our model and mice. Mice have a twelve to fifteen day remodeling cycle, but our model has a 100-day remodeling cycle for humans^37^. We held constant the number of remodeling cycles between mice and our model when comparing them. We assumed that the two month old mice could be modeled as having undergone four cycles, the three month old mice could be modeled as having undergone six cycles, and that the six month old mice could be modeled as having undergone twelve cycles. Thus, we ran the mathematical model for either four, six, or twelve 100-day cycles.

### Wnt-10b fold change

For our model, we take fold change of Wnt-10b to be

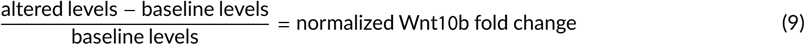

Note that the notation for “altered” and “baseline” remains consistent with the previous section except Wnt-10b is in place of BV/TV. This is done so that the model will produce a balanced remodeling cycle (returning to initial steady state) when *Wnt* = 0, representing a normal baseline level of Wnt-10b.

### Numerical methods

The system of ODEs for the populations of the bone cells and the bone volume in Eqs. 1–2 and 4–6 are solved simultaneously using the `ode15s` function in MATLAB, which is a variable-step, variable-order solver based on numerical differentiation formulas of orders one to five for stiff problems. The relative tolerance is set to 10^−7^. The time step is set to 551 steps per 100-day cycle to ensure fine resolution of the peaks of the trajectories. The initial conditions are provided in Table 4.

**TABLE 4.** Initial conditions for the species tracked in Equations 1–2 and 4–6.

| Symbol | Initial condition | Definition | Units |
| --- | --- | --- | --- |
| *S* | 180 | Osteocyte population at time *t* | cells |
| *P* | 0 | Pre-osteoblast population at time *t* | cells |
| *B* | 0 | Osteoblast population at time *t* | cells |
| *C* | 0 | Osteoclast population at time *t* | cells |
| *z* | 100 | Relative bone volume at time *t* | % |

### Parameter estimation

We fit the parameters *α*_3*adj*_, *β*_1*adj*_, *β*_2*adj*_, and *K*_*M*_ using the MATLAB nonlinear least-squares function `lsqcurvefit`. The multicycle effect of Wnt-10b is on bone volume. Thus, we solve the system of ODEs using the numerical methods described in the previous section for multiple cycles. The ODE solver is called for one remodeling cycle at a time. The solutions from the final time at the end of one cycle are used as the initial conditions for the subsequent cycle with two exceptions: (1) cell populations that have a value less than one are set to zero to not allow fractional values less than one discrete cell to persist, and (2) the osteocyte population is reset to 180 cells at the start of each cycle. The process is repeated for an integer number of cycles. The bone volume calculated at the end of the series of remodeling cycles is taken as the model output for the parameter estimation algorithm and is compared to the normalized BV/TV data. The input is the magnitude of normalized Wnt-10b fold change. We assume that the saturation parameter *K*_*M*_ could take on a value between 1 and 100, and we bound the parameter fitting accordingly. We bound *β*_1*adj*_ between zero and one based on the magnitudes of the other parameters. We set the value of

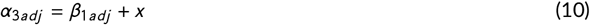

where 0 < x ≤ 1 is an estimated parameter to make *α*_3*adj*_ > *β*_1*adj*_ to ensure that the total effect of increasing Wnt-10b on pre-osteoblasts yields a net increase in formation. Since *β*_2*adj*_ and *β*_2_ collectively impact osteoblast apoptosis, we check that the net apoptosis rate satisfies

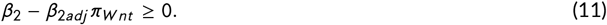

For `lsqcurvefit` the optimization options are set to have step tolerance of 10^−14^, function tolerance of 10^−14^, optimality tolerance of 10^−6^, and 10,000 maximum function evaluations.

To explore the parameter space, we ran `lsqcurvefit` utilizing a multistart algorithm, sampling initial parameter guesses first via random number generation using `rand`. Using the squared 2-norm of the residuals (`resnorm`) to sort the resulting best fit parameters, we narrowed the sampling space and repeating the multistart algorithm using `lhsdesign` for latin hypercube sampling of the initial parameter guesses. We compiled all of the multistart `lsqcurvefit` results and sorted them by increasing `resnorm`. We reduced our parameter set by cutting the parameter sets that produced results that fell outside of 10 percent of the smallest (best fit) `resnorm` of 261.9. We ignored any results with a parameter value less than 1 × 10^−4^ because Wnt-10b should have an effect on these parameters (i.e., we want them to be large enough compared to the other parameters we did not adjust). Additionally, we eliminate parameter sets that resulted when the parameter estimation routine stopped because the upper bound of any parameter was reached. From this parameter set, we found that any sets where *x* > 0.1 led to pre-osteoblast production after termination of the production signal from *S*, even when *C* = 0. When the signal is terminated, the pre-osteoblast population *P* should not increase at the end of the cycle. At this point, the rate of change in *P* is only dependent on the last three terms in Equation 4, which simplifies to

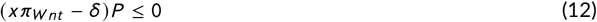

So, we considered as valid only the parameter sets where *x* ≤ *δ* as *π*_*Wnt*_ → 1. We took the average and SD of these results and did a final multistart parameter estimation using MATLAB `normrnd` to sample the space around the parameter averages at finer resolution. Finally, we narrowed our results to be within 1 percent of our new lowest `resnorm` of 277.2. Note: this `resnorm` value is larger than that mentioned earlier for the broader parameter sets that were considered before applying the constraint of Equation 12 and the minimum parameter values of 1 × 10^−4^. We selected our parameters to be the average of this final set of 20 estimated parameter groups. The average and SD for each parameter are listed in Table 5. We noted that the SD of *β*_2*adj*_ was higher than expected, but we found that the model only changed by 2 percent or less when *β*_2*adj*_ is set to the extremes of the SD.

**TABLE 5.** Estimated parameter values and definitions.

| Parameter | Definition | Value | SD | Units |
| --- | --- | --- | --- | --- |
| $\alpha_{3adj}$ | Wnt-10b dependent pre-osteoblast proliferation rate | $2.61 \times 10^{-1}$ | $1.56 \times 10^{-2}$ | day<sup>-1</sup> |
| $\beta_{1adj}$ | Wnt-10b dependent differentiation rate of pre-osteoblasts | $8.33 \times 10^{-2}$ | $1.34 \times 10^{-2}$ | day<sup>-1</sup> |
| $\beta_{2adj}$ | Wnt-10b dependent osteoblast apoptosis rate | $7.10 \times 10^{-4}$ | $7.00 \times 10^{-4}$ | day<sup>-1</sup> |
| $K_M$ | Half saturation for Wnt-10b | 6.26 | $4.78 \times 10^{-2}$ | dimensionless |

### Code sharing

To enable code reuse, we share our MATLAB code including parameter values, scripts for plotting, and documentation in an open-source software repository ^38^. We also include the systems biology markup language (SBML) version of our model for others to reuse readily. This version can be easily converted to Python^39^.

## Results and Discussion

Parameter estimation resulted in parameters (Table 5) that fit the data ^16,17^ within the uncertainty of the altered mice data. Figure 2A shows the results at the end of six or twelve remodeling cycles using the estimated parameters. Figure 2B shows the cumulative changes in bone volume after these remodeling cycles are plotted against varying fold changes in Wnt-10b for four, six, or twelve remodeling cycles. Note that the data points are the same for A and B but are represented on different axes. The parameter fitting results visually agree with the data well. With fitted parameters, the model gives the bone volume progression over multiple remodeling cycles for the full range of Wnt-10b perturbations explored. The initial conditions for these simulations are provided in Table 4. As Wnt-10b increases, the bone volume also increases.

**FIGURE 2.**
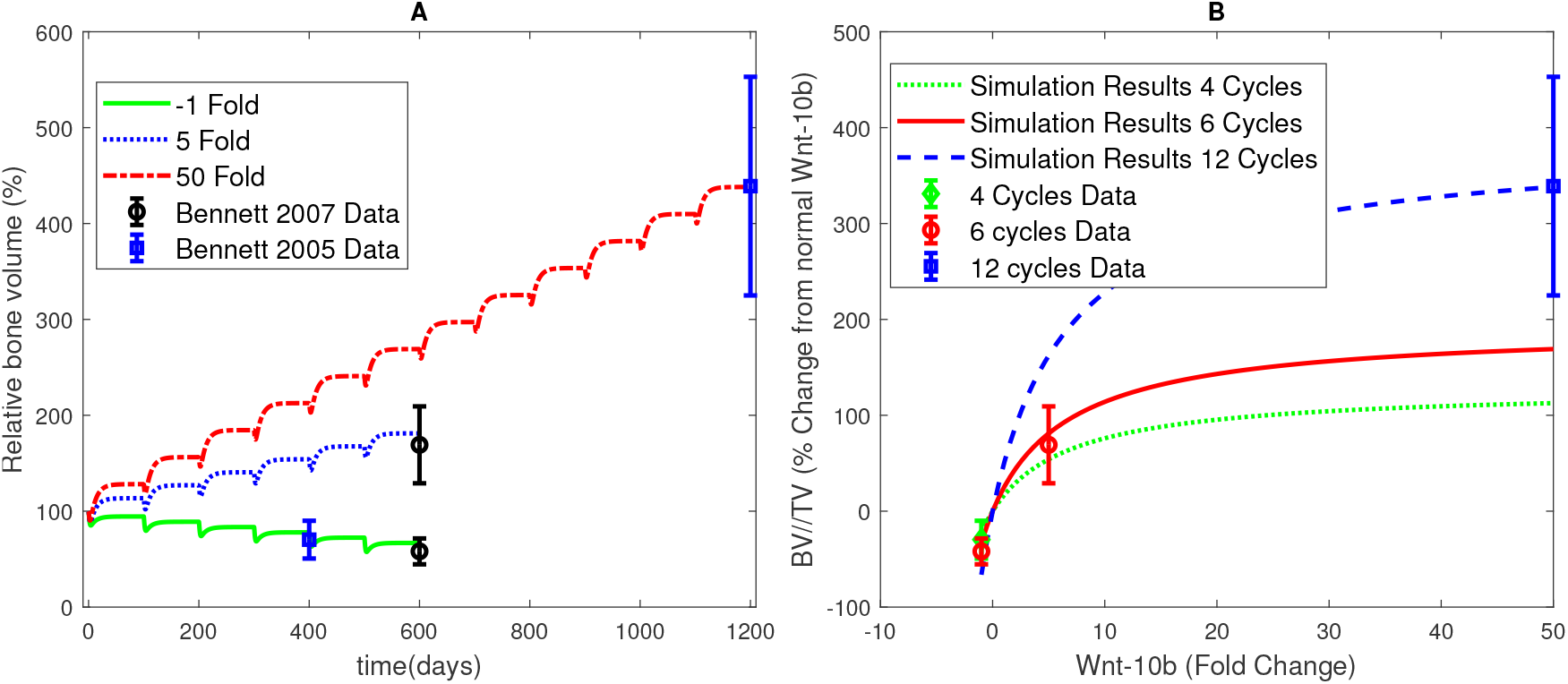
Fitting and simulation results of bone volume and Wnt-10b. A) Fitting results for three normalized Wnt-10b fold changes shown in the legend. These specific fold changes were used to parameterize the model by comparison of the final bone volume with the corresponding data points for the Bennett 2005 data^17^ and the Bennett 2007 data^16^. B) Simulated normalized BV/TV relationship with normalized Wnt-10b fold change. The parameter fitting simulation results produce an acceptable visual fit when compared to the error. Data points were compared with the cumulative final simulated BV/TV after four remodeling cycles (green dotted curve), six remodeling cycles (red solid curve), or twelve remodeling cycles (blue dashed curve). Fold changes of Wnt-10b range from −1 to 50 in this simulation. Note that the data used in A and B is the same but represented on different axes.

To determine if the model is a good predictor of the change in bone volume that occurs when Wnt-10b levels are perturbed, we utilized a separate set of data ^18^. The simulation results fall within the error of the data at six remodeling cycles, but overestimate the results at twelve remodeling cycles (Figure 3). This model validation is particularly good for six or fewer remodeling cycles. The shaded area and the error bar of the data at twelve cycles do overlap to a reasonable extent. The offset at later time points could be due to the fact that the mice in the study used for validation had intermittent doses of Wnt-10b via weekly injections, which possibly allowed for fluctuating Wnt-10b levels. On the contrary, the mice were genetically altered in the studies we used for modeling fitting. Thus, the Wnt-10b levels were presumably constant for the genetically altered mice, and the reported levels of Wnt-10b in the injection experiments may not have captured the dynamics. The shaded area in Figure 3 shows the simulation results within the standard deviation of the Wnt-10b reported fold change, assuming that the fold change is constant throughout the duration of the twelve cycles. The bone volume would shift lower if the Wnt-10b levels were to be allowed to diminish between injections. Alternatively, the offset could show a decreased sensitivity to Wnt-10b over long term treatment.

**FIGURE 3.**
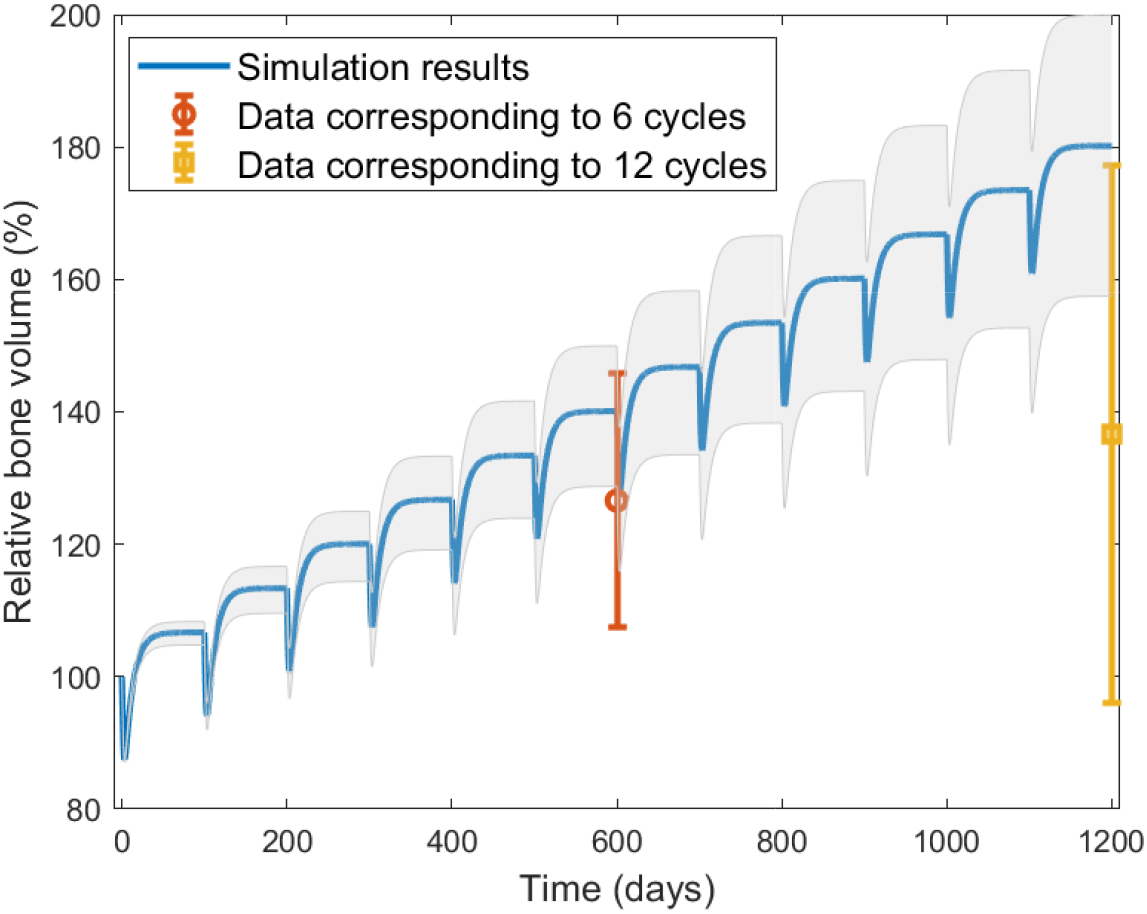
Validation of model with data from Roser-Page et al.^18^. The simulation shown is the result of running the model with a 1.8 normalized fold change of Wnt-10b for twelve remodeling cycles. The grey shaded area shows the simulation results for the standard deviation (± 0.6) of the reported Wnt-10b fold change. The data^18^ corresponds to six and twelve remodeling cycles. The simulation falls within the error of the data for six remodeling cycles, but overestimates for twelve remodeling cycles.

For each fold change in Wnt-10b shown in Figure 2A, we also determined the corresponding numbers of the activated populations of osteocytes, pre-osteoblasts, osteoblasts, and osteoclasts (Figure 4). The results show a positive correlation with pre-osteoblast and osteoblast populations and a slight negative correlation with osteoclasts (Figure 5). Even though we did not alter the osteoclast equation (Equation 2), the population is related to the ratio of pre-osteoblasts to osteoblasts. The model predicts that the ratio of pre-osteoblasts to osteoblasts and the ratio of osteoclasts to osteoblasts both decrease with increasing Wnt-10b (Figure 6). The cumulative numbers of cells of each type were determined by integrating the areas under the curves (AUC) shown in Figure 5 using the `trapz` function in MATLAB. The AUC were used for comparison instead of the dynamic number of cells given that the initial conditions for numbers of activated cells of all three types are zero.

**FIGURE 4.**
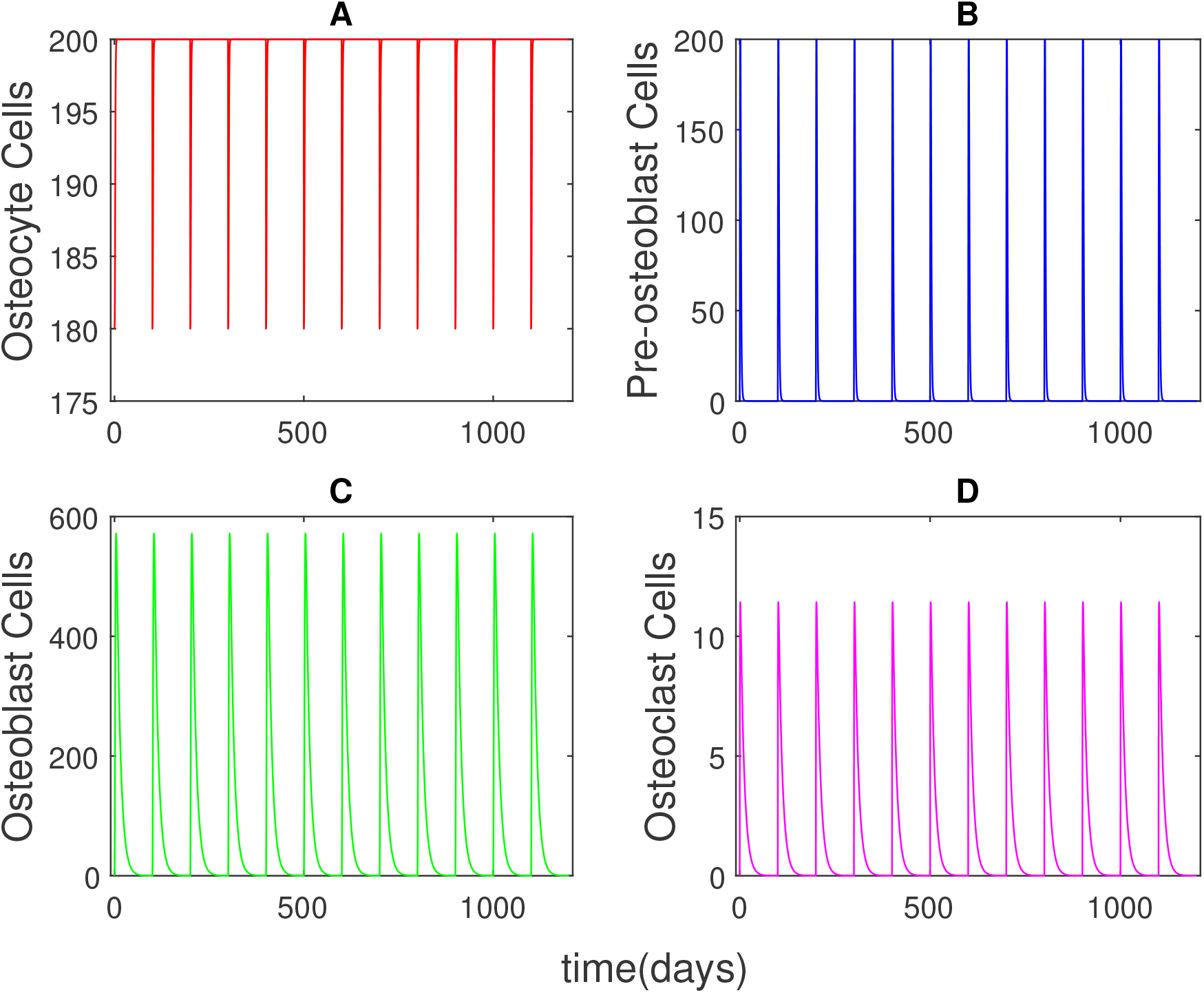
Activated cell population dynamic results for (A) osteocytes, (B) pre-osteoblasts, (C) osteoblasts, and (D) osteoclasts at a normalized 50-fold increase in Wnt-10b. The activated cell populations follow identical dynamics for each remodeling cycle while Wnt-10b concentration is constant.

**FIGURE 5.**
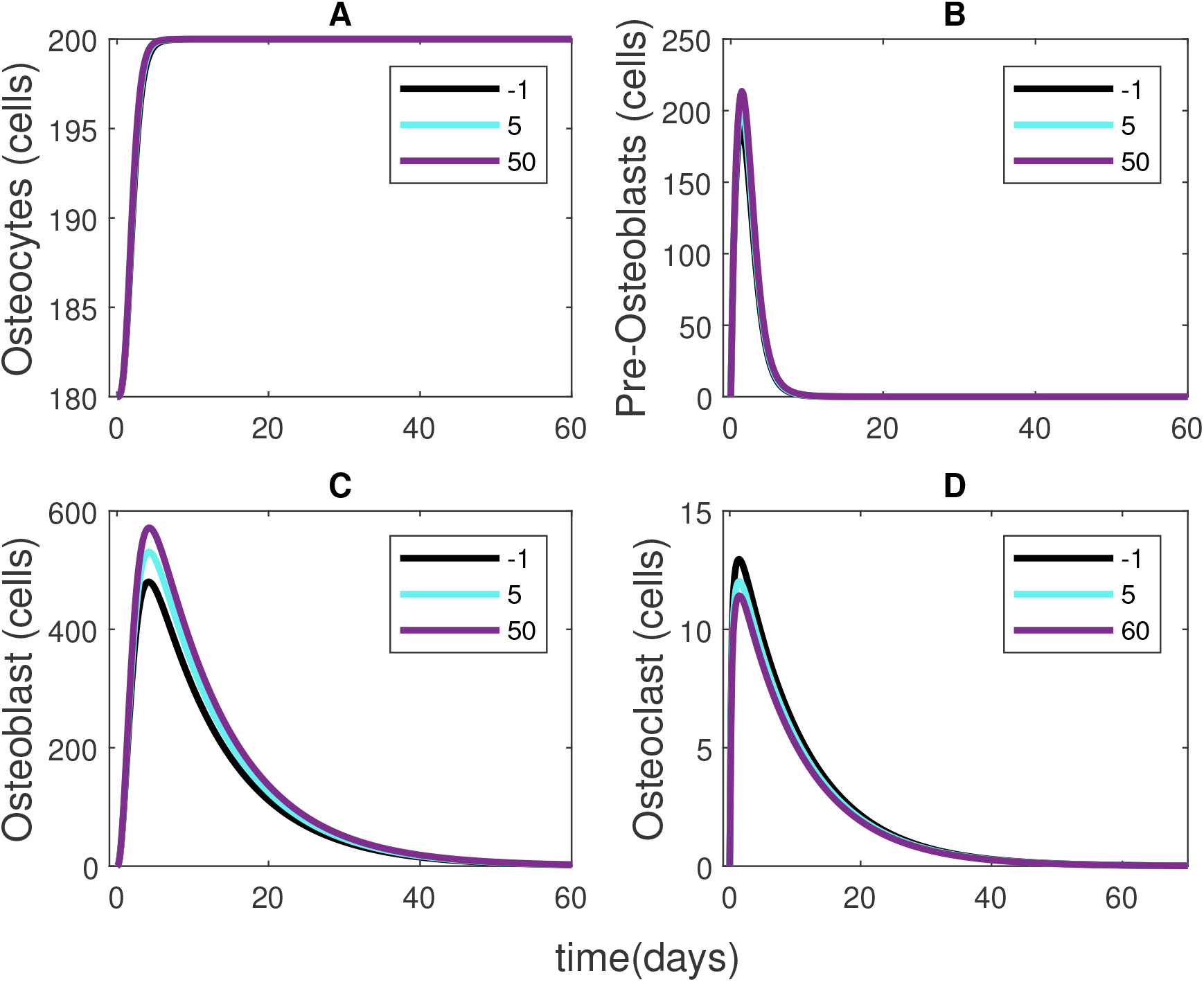
Activated cell population results for a single remodeling cycle with varying Wnt-10b concentrations shown in the legends. (A) The osteocyte dynamics show little change across Wnt-10b concentration. (B) The pre-osteoblast dynamics show a very slight positive correlation with Wnt-10b. (C) The osteoblast dynamics show a positive correlation with Wnt-10b. (D) Osteoclasts decrease with increasing Wnt-10b. Note that the cycle continues to 100 days with no further changes in the cell populations.

**FIGURE 6.**
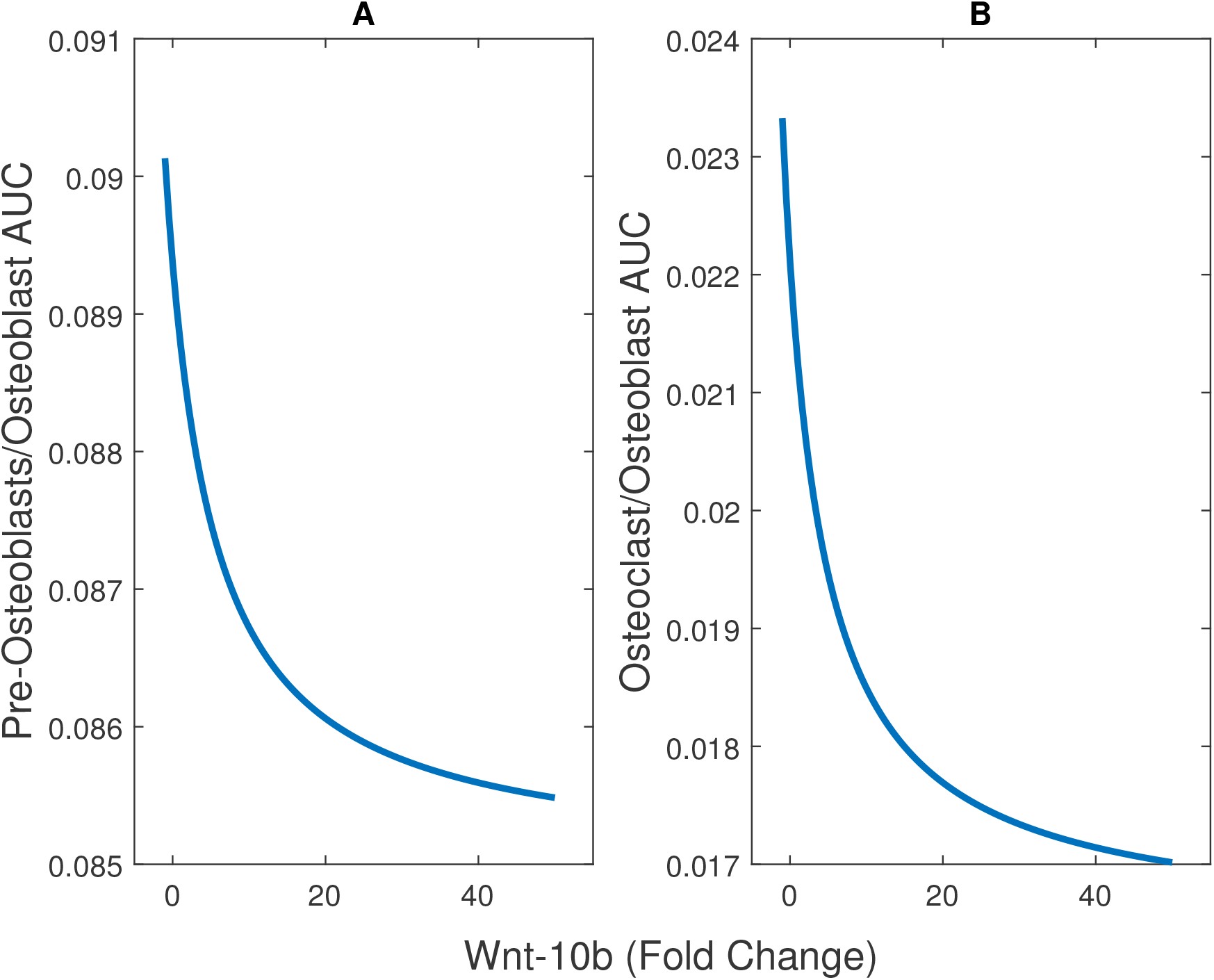
Ratios of the cumulative numbers (area under the curve, AUC) of (A) pre-osteoblasts and (B) osteoclasts to osteoblasts. As Wnt-10b increases, both of the ratios decrease.

The reduction of osteoclasts population has been experimentally tested for mice with a normalized 5-fold increase in Wnt-10b, and the change in the number of osteoclasts on the perimeter of the bone was not statistically significant ^16^. However, based on the physiology of bone remodeling, it is expected that a change in the ratio of pre-osteoblast and osteoblasts would alter osteoclast populations through the RANK/RANKL/OPG pathway^27,40,41^. In this pathway, pre-osteoblasts, osteoblasts, and osteocytes secrete RANKL that binds to RANK receptors on osteoclasts thus increasing osteoclastogenesis. Osteoblasts and osteocytes also secrete OPG, which is a competitive inhibitor of RANKL (Figure 7). As the osteocyte dynamics are shown to be relatively consistent, OPG levels change due to a change in osteoblast dynamics. It is also important to note that the predicted change in osteoclast population is small enough that it would be hard to quantify experimentally. Determining the indirect relationship between activated osteoclast population and Wnt-10b would help to further validate the osteoclast population aspect of the model.

**FIGURE 7.**
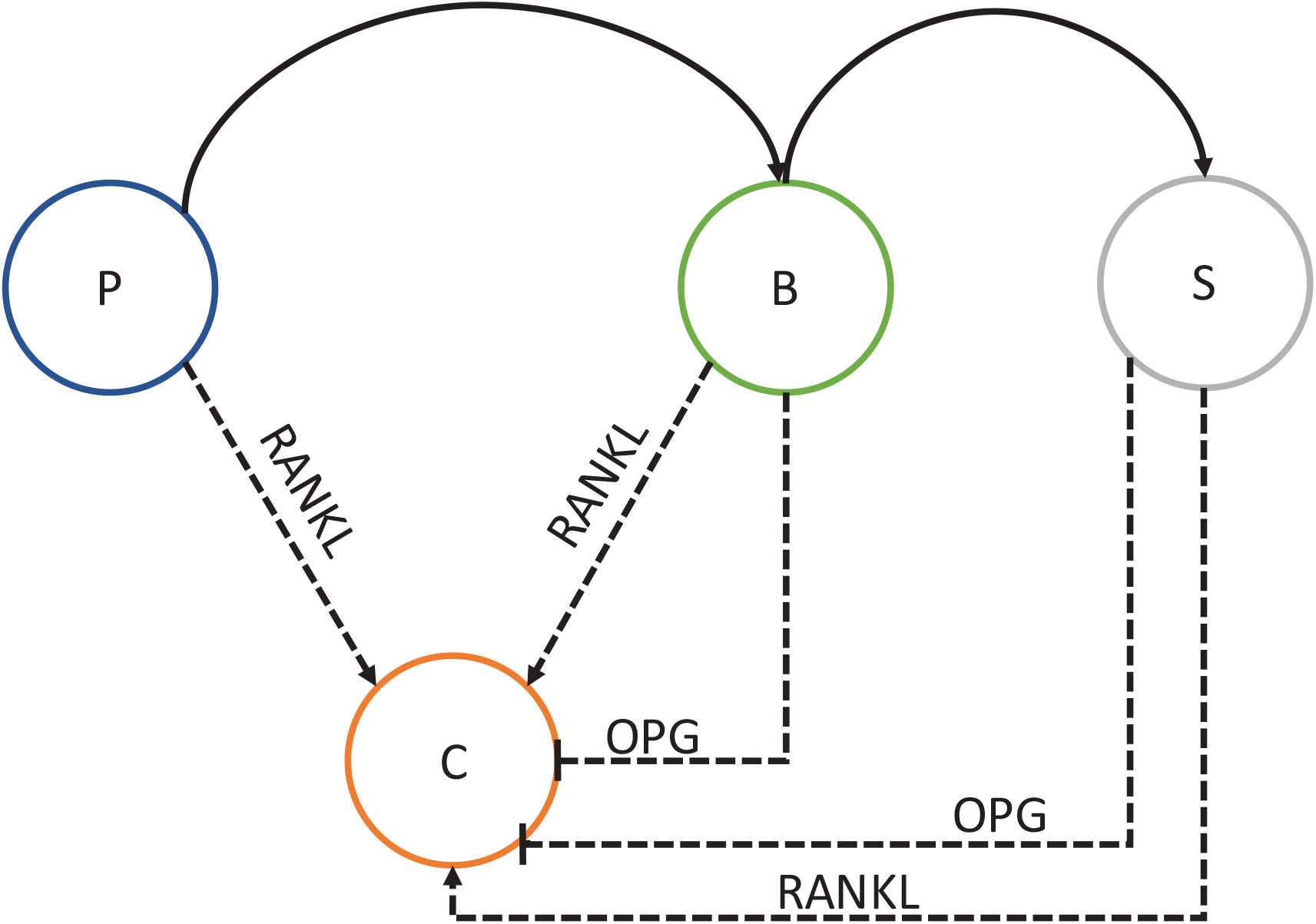
RANK/RANKL/OPG pathway where osteoblastic cell populations regulate the osteoclasts (C) cell populations. Pre-osteoblasts (P), osteoblasts (B), and osteocytes(S) secrete RANKL causing an increase in osteoclasts. Osteoblasts and osteocytes also inhibit osteoclastogenesis by secreting OPG, a competitive inhibitor of RANKL.

This model does provide insight into bone metabolism, but it is not without its limitations. The data utilized was normalized mice data, and the remodeling cycle was set to 100 days based on the Graham 2013 model instead of a typical 200 day human remodeling time. We also do not account for the variation of data within a population. These issues can be resolved if more data becomes available. In the future this model could be expanded to include other direct relationships with cell signaling molecules of interest such as TNF-*α* (experimentally shown to alter osteoclast formation and activity) and IL-6 (experimentally shown to alter osteoclast formation)^42^.

## Conclusions

Understanding how Wnt-10b alters the dynamics of bone metabolism could lead to new therapeutic targets for osteoporosis. Though there are a few published models focused on bone metabolism, the model developed here provides new insight on how chronic changes in Wnt-10b affect a single bone remodeling cycle or a sequence of cycles. Experimentally Wnt-10b has been shown to increase osteoblastogenesis and decrease osteoblast apoptosis rates. Our model is able to generate the expected changes in pre-osteoblasts and osteoblasts and is able to predict changes in bone volume caused by Wnt-10b. Our model also points to a slight decrease in osteoclasts that is hard to determine experimentally but is mechanistically sound. These results could be used to design future experiments that could further enhance understanding of how Wnt-10b regulates bone metabolism.

## Abbreviations

AUC: area under the curve
BMU: basic multicellular units
BV/TV: trabecular bone volume per total volume
ODEs: ordinary differential equations
PTH: parathyroid hormone
RANK: receptor activator of nuclear factor kappa B
RANKL: receptor activator of nuclear factor kappa B ligand
OPG: osteoprotegerin
SBML: systems biology markup language
SD: standard deviation
Wnt: winglessrelated integration site family

## Acknowledgments

Research reported in this publication was supported by the National Institute of General Medical Sciences of the National Institutes of Health under award number R35GM133763. The content is solely the responsibility of the authors and does not necessarily represent the official views of the National Institutes of Health.

We would like to thank Dr. Joshua D. Ramsey at Oklahoma State University for the guidance provided as a master’s committee member for C. V. Cook. We would also like to thank Dr. Stelios T. Andreadis, Dr. Viviana Monje-Galvan, and Dr. Rudiyanto Gunawan at the University at Buffalo for their suggestions provided as a Ph.D. committee members for C. V. Cook. Lastly, we would like to thank the other members of the Systems Biomedicine & Pharmaceutics Lab at the University at Buffalo for providing feedback on this project.

